# Walking pattern analysis using deep learning for energy harvesting smart shoes with IoT

**DOI:** 10.1101/2020.05.10.087197

**Authors:** Neel Shah, Laxit Kamdar, Drashti Gokalgandhi, Ninad Mehendale

## Abstract

Wearable Health Devices (WHDs) benefit people to monitor their health status and have become a necessity in today’s world. The smart shoe is the type of WHD, that provides comfort, convenience, and fitness tracking. Hence smart shoes can be considered as one of the most useful innovations in the field of wearable devices. In this paper, we propose a unique system, in which the smart shoes are capable of energy harvesting when the user is walking, running, dancing, or carrying out any other similar activities. This generated power can be used to charge portable devices (like mobile) and to light up the LED torch. It also has Wi-Fi-that allows it to get connected to smartphones or any device on a cloud. The recorded data was used to determine the walking pattern of the user (gait analysis) using deep learning. The overall classification accuracy obtained with proposed smart shoes could reach up to 96.2 %. This gait analysis can be further used for detecting any injury or disorder that the shoe user is suffering from. One more unique feature of the proposed smart shoe is its capability of adjusting the size by using inflatable technology as per the user’s comfort.

## 1 Introduction

Consumer dependency on wearable devices has grown rapidly in the past few decades[1]. As the wearable devices are now capable of monitoring the health of a person, they are now termed as Wearable Health Devices or WHDs. These devices provide data, wherein the device sends out suggestions on how one can improvise in their lifestyle for healthful living. Many devices like smartwatches, smart eyewear, fitness tracker, smart clothing are a few examples of WHDs. One such innovation in wearable health devices is smart shoes that are trending and evolving historically in the engineering field. Smart shoe is the technological innovation in the ordinary shoe with the high tech features that record biometric data and activities of the user. This data is recorded and transferred via Bluetooth or Wi-Fi, to mobile applications or computers for the analysis purpose. Devices like sensors, controllers, wireless devices, accelerometer [2], gyroscope, and magnetometer [3] are used for understanding the walking pattern and collecting data for the gait analysis [4]. As smart shoes are capable of monitoring the physical health and to offer suggestions for improvement in the wearer’s lifestyle, there is a need and demand for it are increasing day by day.

In this paper, along with keeping the track of user’s health using IoT, harvesting electricity by walking, running, or carrying out other similar activities has been demonstrated. The power generation is done by using piezoelectric plates, which are placed at the base of the sole. As the user starts a normal activity such as walking or running, the pressure is applied on the shoe, utilizing this pressure, energy is harvested by the piezoelectric sensors that can be used to charge mobile phones, Bluetooth earphones, or other such similar devices. Figure 1 shows the proposed diagram of smart shoes. Energy is harvested, when a person starts walking or running. A maximum output of 9 volts is generated while running using these piezoelectric plates. Piezoelectric plates generate ac voltage and that is converted to dc voltage by using rectifier. The dc voltage is given to the voltage regulator. The output of regulator is given to charge the battery. This battery provides power to LED via a switch, USB connector for external charging, and to the NodeMCU microcontroller. Accelerometer and gyroscopic sensor is interfaced to NodeMCU.The data is generated with the help of MPU6050 sensor and it is transferred to ThingSpeak (cloud storage) via WiFI of NodeMCU. The sensor data is then displayed on the channel. The data was processed further for Gait analysis using deep learning. Another distinct feature of smart shoes is electronic auto-fitting using inflatable sole. L293d motor driver IC is used for driving the air pump motor. The air pump motor is used to inflate the tube according to the remaining space in smart shoes for comfort auto-fitting.

**Figure 1:**
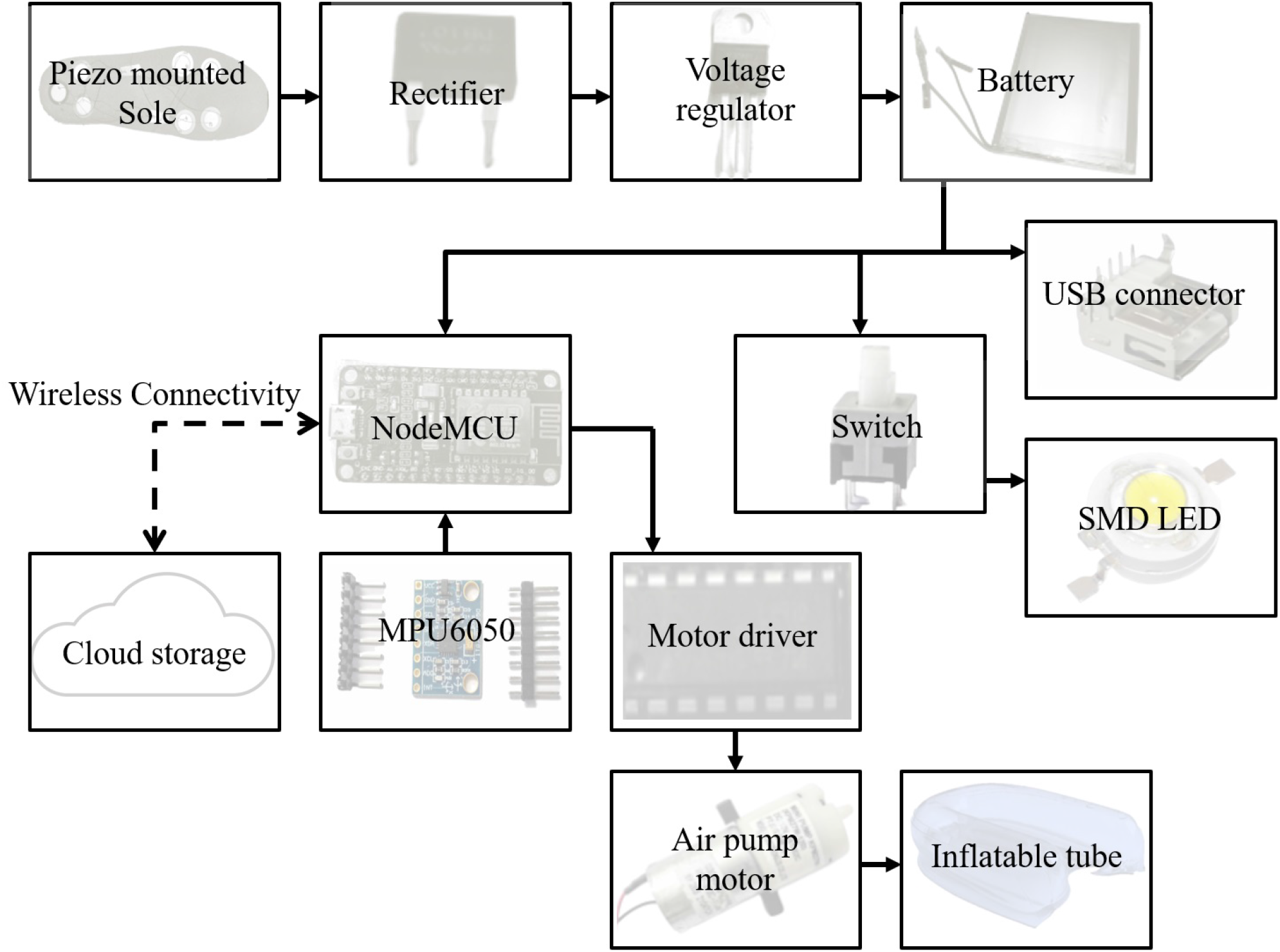
Block diagram of Energy harvesting smart shoes with IoT.The energy harvested from the piezoelectric crystals is stored in the battery, that is used to power up the NodeMCU, torch, and USB charging port. NodeMCU data obtained via MPU6050 was used to analyse the walking pattern of the user. An inflatable tube and air pump are used for achieving comfort fitting.

## 2 Literature review

At present many smart shoes have been developed[5, 6, 7]. In general, there are three main types of smart shoes: Electronic based smart shoes[8, 5], Mechanical based smart shoes, and Electro mechanical smart shoes. Electronic smart shoes can have micro-controller and sensors, wireless connectivity, and navigation system [9] inside it. There are multiple shoes reported in the literature for visually-impaired [10] or physically challenged people. Bluetooth enabled smart shoes [11] are also developed. A big brand like Adidas [12] has developed smart shoes with intelligent cushioning with the help of magnet and sensor. The present manuscript takes advantage of all these reported technologies and further adds IoT functionality to transfers the data on cloud storage i.e on ThingSpeak. The data was analyzed using deep learning for gait analysis. Mechanical smart shoes are passive pump based smart shoes. The passive pump [13] technology allows customization of the shape of each shoe to fit the precise shape of each foot. The reported smart shoe has integrated an inflatable urethane bladder with an air compressor at the side of the shoe and a pressure release valve on the heel. All the technologies reported in the literature, has a manual air compressor, where user efforts of pressing the button consecutively are required. On the other hand, the shoes we designed can be adjusted electronically with a single press of a button, tubes are inflated automatically. Similar technology was reported by VectraSense smart shoes [14]. VectraSense presented a new computerized shoe named ‘Verb for Shoe’. This footwear was skilled for sensing the user’s activity level and automatically adjusting itself to improve comfort and performance. The shoes were integrated with an embedded computer that could learn the patterns and adjust the fitting according to the comfort of the user via the air bladder technology. The electromechanical smart shoes are of two types. The first one is the generator embedded into the smart shoes. These generators could be electromagnetic, piezoelectric, or solar panel-based [15]. The second type of electro-mechanical smart shoe is motors induced inside a smart shoe to give a mechanical vibration. The reported generator embedded smart shoe has a solar panel generator along with a piezoelectric generator. The energy generated is used to charge the smart shoe battery. However, recently reported an electromagnetic generator and a piezoelectric generator [16, 17] are a more promising approach for energy harvesting as they can generate high output current level and voltage. With this consideration, we have used piezoelectric plates for the generation of power in our smart shoes. The power generated can even be used as a power bank to charge the other devices. The motor induced [6] inside a smart shoe is also reported in the literature. This motor is used for training foot progression angle during walking for the patients with osteoarthritis. There are some recent reports that use deep learning algorithms to determine the walking problem or walking pattern. Deep learning can also be used for classification [18, 19, 20].

## 3 Methodology

### 3.1 Hardware

NodeMcu was acquired from RS Electronics. MPU6050 sensor was purchased online from the element14 website. Air pump motor was acquired from robu.in which was used to inflate tubings inside sole, the tubings bought locally. Motor driver IC L293D was also acquired from robu.in and it was used to drive the air pump motor. SMD LED (1 watt) was purchased from electronicscomp and used as a torchlight. Piezoelectric plates were purchased locally from Visha kits. USB connector was also purchased locally from Shanti electronics. Regular shoes were brought from Nike showroom of size 12. Battery and other small components such as copper-clad, FeCL3, etc. were purchased from Vega robokits. Insole and sponge were bought locally.

### 3.2 Softwares

Arduino IDE free software version 1.8.4 was used. Fritzing version 0.87 open source was used to design the PCB. ThingSpeak free web service was used to transfer data over the internet. MATLAB version 19b was used along with the Machine learning toolbox.

### 3.3 Shoes fabrication

Features we considered while designing the smart shoes were energy harvesting, LED as a torchlight, power bank, pedometer, walking pattern analysis (gait analysis), and auto-fitting (comfort using inflatable tube). Considering these features the circuit designed with the help of datasheets and interfacing diagrams. After designing the circuit, all the components were acquired. PCB designing is being done using Fritzing software. Figure 2 (a) shows the connection diagram of the components mounted on the breadboard. These connections were done as per the datasheets. Figure 2 (b) shows the schematic diagram of the smart shoes. The schematic diagram consists of an MPU6050 sensor interfaced with the NodeMCU. Motor driver L293d interfacing with NodeMCU is also shown in fig. 2 (b). Input to DB107 bridge rectifier IC is given from the piezoelectric sensor. IC7805 is used for voltage regulation. SPDT switch is used to turn on or off the LED torch. The USB connector is attached for the charging of devices. Figure 2 (c) shows the PCB layout of the designed PCB for smart shoes. PCB was designed using the Fritzing software. And fabricated using standard negative photolithography and FeCl3 etching process. Figure 3 (a) shows the inner sole of the smart shoes. Plain rubber inner sole is converted into energy harvesting sole. Energy is harvested using piezoelectric plates. Maximum 9 volts is generated from the piezoelectric plate and to increase the output current 7 piezoelectric plates were connected parallelly. The electrical charge generated by applying mechanical stress on the piezoelectric sensor mounted on the inner sole was being converted to dc voltage with the help of the DB107 bridge rectifier. The converted dc voltage is further given to IC 7805. 7805 is a voltage regulator. It constantly gave 5V output whenever the input dc voltage was more than 5.7 volts. The output of 7805 is further given to charge the battery. The battery-powered the circuitry. SMD LED was being turned on or off using a Single pole double throw (SPDT) switch. SMD LED was used as a torchlight in darkness or in the night. We were also able to charge the mobile using a USB cable plugged to the USB connector in the front of the shoes. MPU6050 sensor was interfaced with the NodeMCU microcontroller. The data was being generated with the help of the MPU6050 sensor. Figure 3 (b) shows the final PCB of smart shoes fabricated and components are mounted using a standard soldering technique. PCB is designed using Fritzing software. Figure 3 (c) shows the assembly of smart shoes. The PCB is mounted directly inside the main bottom sole of smart shoes. The battery is attached to the PCB using a switch. The layer of sponge is placed in the remaining space of the main sole for protecting PCB from any damage. The inner sole with inflatable tubing is then sandwiched with the piezoelectric sole and placed on the top facing downwards. Figure 3 (d) shows the final fabricated smart shoes. It is ready for testing and using. There is a slot on the front side of the smart shoe for LED and USB charging.

**Figure 2:**
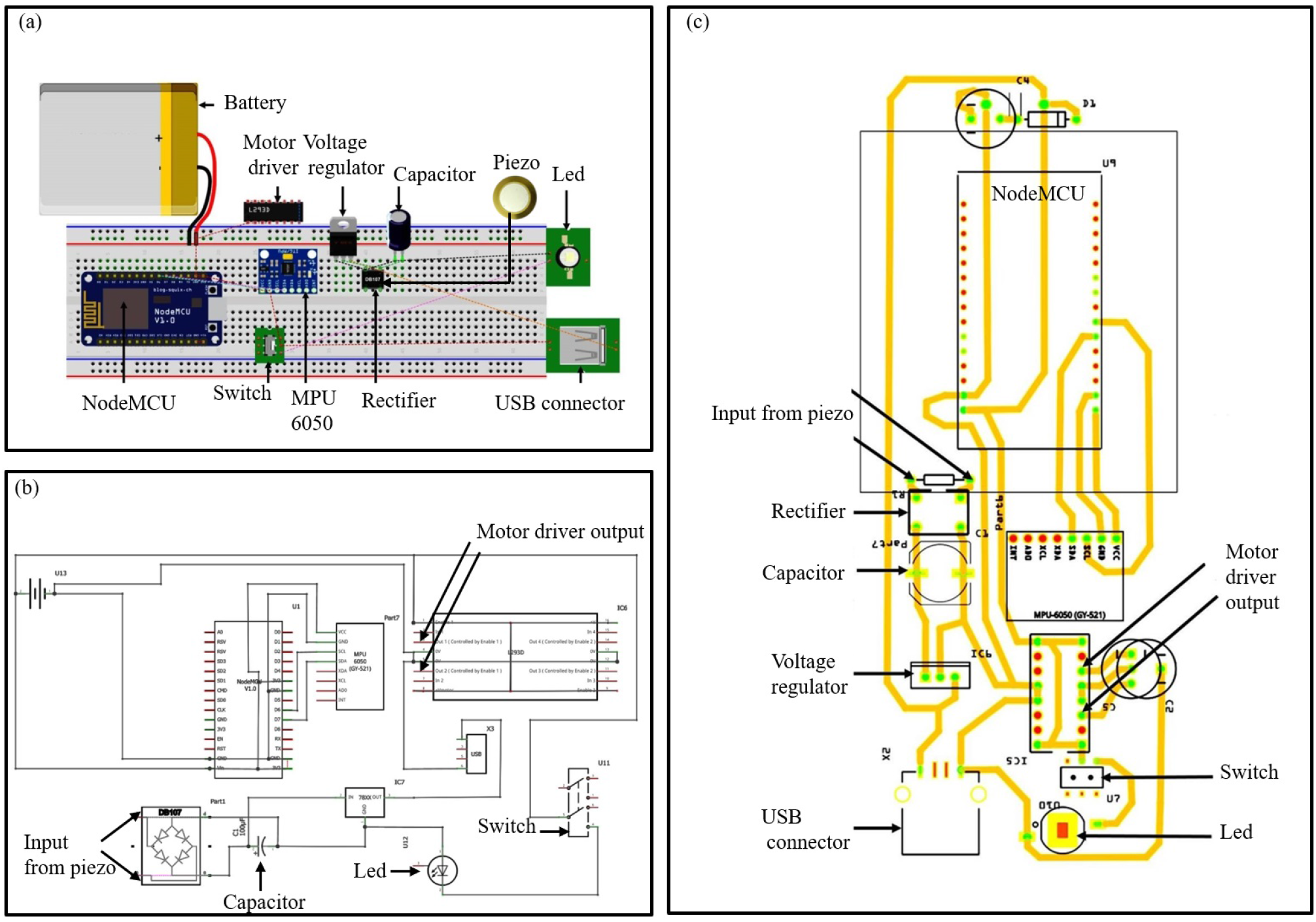
PCB designing procedure of smart shoes using Fritzing. (a) Connection diagram of smart shoes on a breadboard using Fritzing. (b) Schematic diagram of smart shoes. (c) Final designed PCB layout of smart shoes.

**Figure 3:**
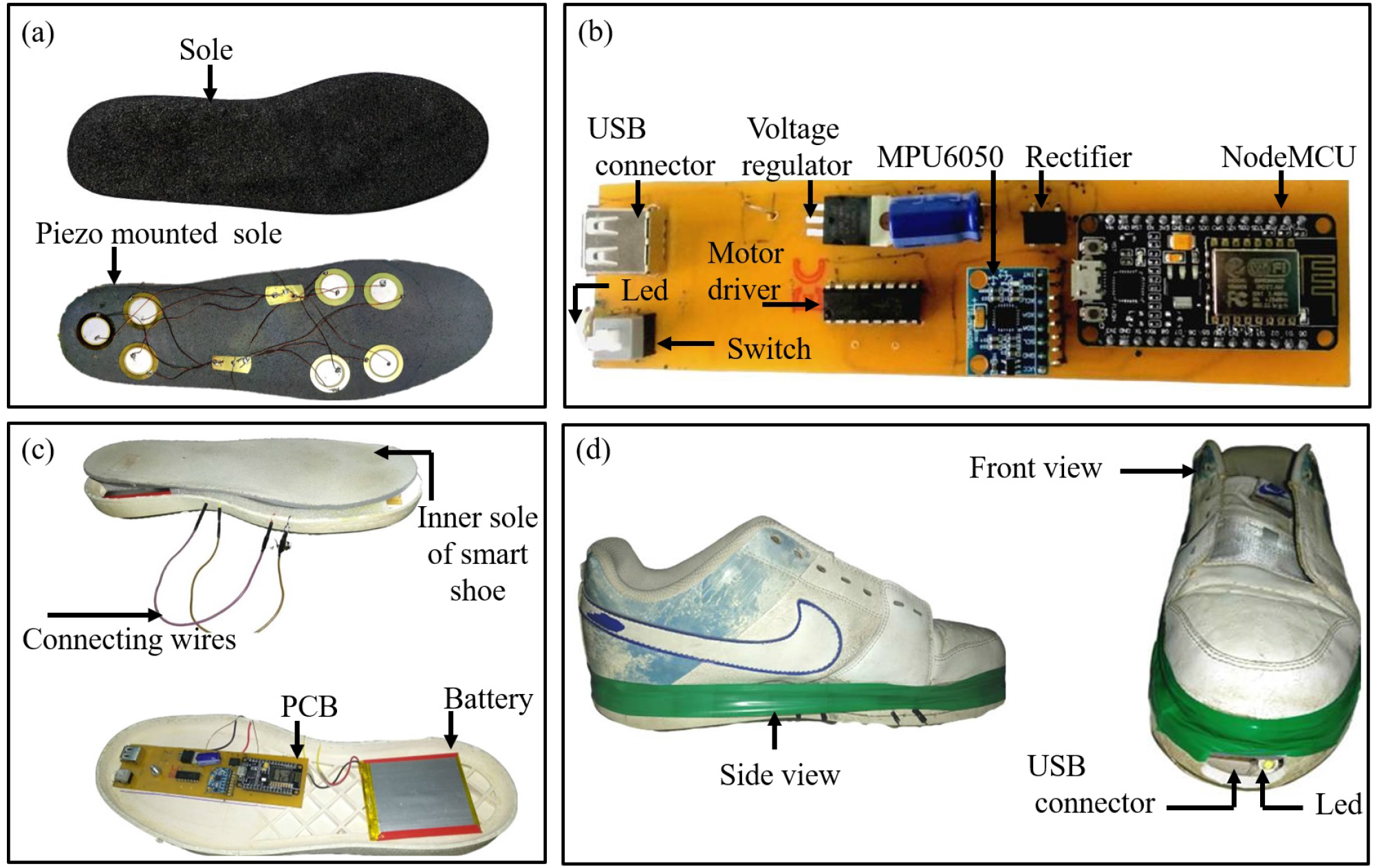
Fabrication procedure of smart shoes (a) Inner rubber sole converted to power generation sole using piezo. (b) Picture of fabricated PCB of smart shoes. (c) Fabrication of outer sole with the PCB, battery, and Inner sole placed on it. (d) Final fabricated smart shoes with inflatable sole and a slot for LED and USB charging in front.

### 3.4 Microcontroller programming

For initial testing with IoT data transfer, Arduino IDE software was used. For serial communication library was used initially for testing. For WiFI communication ESP8266 board package was installed from the board manager. The code for MPU6050 sensor was developed using embedded C (refer supplementary informatiom S1 for the Code). Also API key (e.g. KWRY5OECU9TVAPAO) of the channel created on ThingSpeak for receiving data from NodeMCU must be mentioned in the code.The Service Set Identifier (SSID) and password for Wi-Fi connection must be integrated into the code before compiling and uploading.

### 3.5 Deep learning algorithm for gait analysis

We have used 13-layer deep learning architecture for performing gait analysis. The input data size of 1×784 obtained from the ThingSpeak channel was updated every second. The first layer is the input layer which used zero crossings and truncation to find periodic patterns in received signals. The second, fifth and eight-layer is a convolutional layer. This layer applies convolutional sliding filters to the data. The third, sixth, and ninth layer is batch normalization. A batch normalization layer standardizes any input channel over a mini-batch. Using the batch normalization, processing time is decreased and it also it reduces the sensitivity to network initialization. The fourth, seventh, and tenth layer is ReLU layer. A Rectified Linear Unit (ReLU) performs a threshold operation for each input variable, where any value that is less than zero is set to zero. The data from the tenth ReLu layer was fed to a fully connected network layer. In the fully connected layers, the input is multiplied with a weight matrix. After multiplying, a bias vector is also added to it. The twelfth layer was the softmax layer. The last layer was the classification layer. This classification layer measures the cross-entropy loss of mutually exclusive groups for 8 class classification problems.

## 4 Results and discussions

Figure 4 (a) shows the plot of accuracy versus the number of iterations. As the number of iterations goes on increasing the accuracy of deep learning classification algorithms was saturated. We stopped after 4 epochs of 40 iterations each. After training the accuracy reached up to 96.2% Figure 4 (b) shows loss versus iteration plot. As the number of iteration and accuracy increases the loss also goes down exponentially. We could achieve loss as low as 0.2 after 160 iterations. Figure 4 (c) is of flowchart explaining the working of the smart shoes. Firstly, the sensor data is generated by the MPU6050 sensor is gathered for further process.

**Figure 4:**
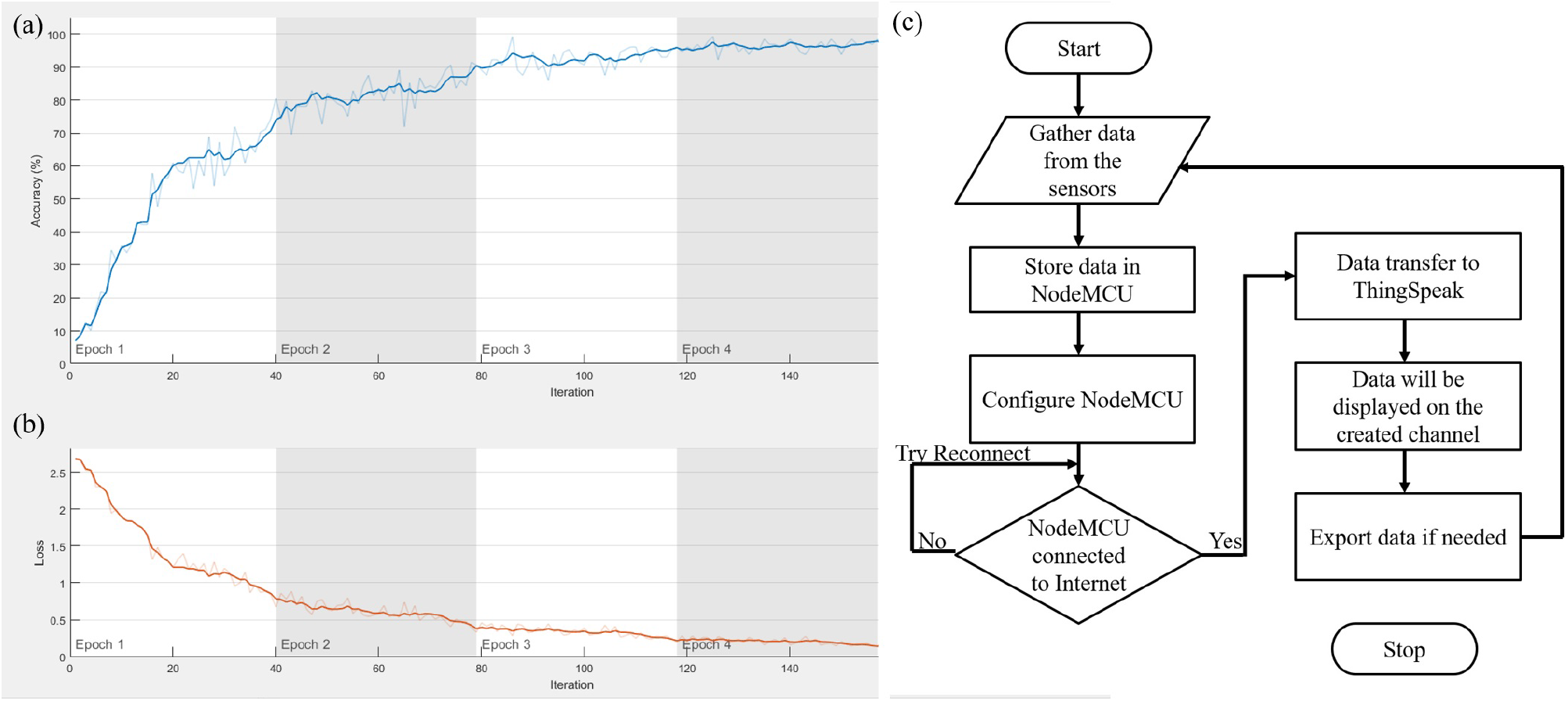
Accuracy (a) and losses (b) plotted against iteration in deep learning networks (c)Flowchart of smart shoe data transmission over IoT using cloud data service ThingSpeak.

This data is temporarily stored in NodeMCU. NodeMCU is programmed and configured for a Wi-Fi connection. If the Internet connection is done successfully, then the data is transferred to ThingSpeak using API write key of the channel i.e on the cloud server. The data transferred from NodeMCU to ThingSpeak will be displayed on the created channel. This data is further exported to MATLAB for further processing. If the NodeMCU is not connected to the Internet then the shoe will keep trying to reconnect until data is transferred.

Classification accuracy is determined by the confusion matrix (fig. 5). There are 8 desired classes and 8 predicted classes. The number of patterns in each desired class was 500 and hence we were expecting all the diagonal elements (i.e. true positive) to be 500. We got a maximum of 498/500 in class 5 of the mid-swing class. whereas minimum accuracy appeared in the foot-flat class. It was expected because Mid-swing has very fewer chances of getting classified in either of the swings (i.e. initial and late) and also the negligible chance getting classified in any of the stance classes. We could figure out for proper reproduction of all classes at least 500 samples per second data was required. We set it to 784 samples per second. Also, FPS of ground truth video was required to be more than 500 frames, we have used 1000 FPS and dropped a few frames from the initial and the final part of the video to match the 784 frames to each sample. 5 such sample ground truths after binarization are attached in supplementary video SV1.

**Figure 5:**
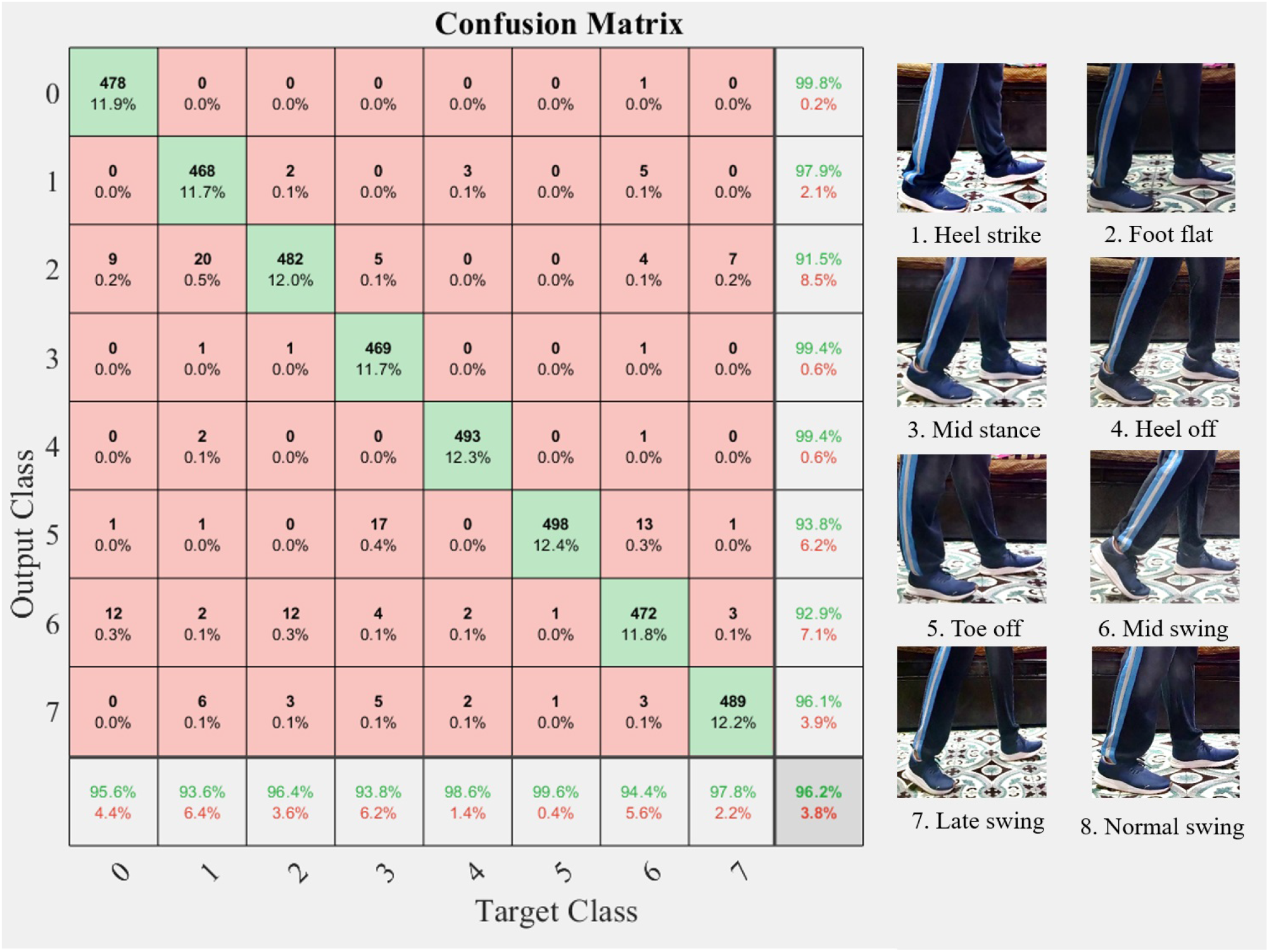
Confusion matrix obtained during the testing phase of smart shoes. The walking pattern is classified into 8 classes, where 5 of them are classified into the stance phase and the remaining 3 are classified into the swing phase. Stance phase includes heel strike, foot flat, mid stance, heel off, and toe-off whereas swing phase has mid-swing, initial swing and late swing.

As shown in figure 5, the walking pattern is divided into 8 classes wherein, five classes are of stance phase and three are swing phase. Each phase represents a gait cycle of the walking pattern as the foot moves or touches the ground. The stance phase includes heel strike, foot flat, mid stance, heel off, and toe-off. The swing phase is further categorized as initial swing, mid-swing, and late swing. The heel strike stage is the first phase of the gait cycle wherein the heel of the first moving foot touches the ground. At the next phase, i.e. the foot flat phase the sole of the moving foot touches the ground. In the next phase i.e. mid stance, the leg of the first moving foot is perpendicular to the ground and momentarily stopped while the other leg moves forward. In the fourth phase, in the heel off stage, the heel begins to move away from the ground and for the fifth stage, in toe-off, the first moving foot leaves the ground while the sole of the other foot touches the ground. In the swing phase, the mid-swing is the step forward with the foot of the ground and the late swing refers to the stage immediately before the next cycle begins. Along with all these standard 7 classes reported in literature, one initial swing class was also defined that corresponds to the initial pattern. The 5 gait phases were repetitive and we needed to add visual recording to find out ground truths for comparison and supervisory training of deep learning network. The camera used was Sony Mark-V with 1000 fps full of slow-motion recording (Refer supplementary video SV1). For generating ground truths for the corresponding signals expert doctors from Ninad’s research Lab and Kaushakya Medical foundation hospital were invited. The study was carried out on human subjects with their written consent.

## 5 Conclusions

Smart shoes will be the next big thing in WHDs. The presented smart shoe could give a pathway for future research. Our smart shoes can generate power through piezoelectric plates and it is stored inside the battery. This solves a big issue of the power bank. The generated power can be further utilized to glow the LED that helps the user to walk in dark. The data is logged by interfacing MPU6050 with NodeMCU and it is sent to the ThingSpeak cloud via Wi-Fi. The deep learning algorithm used for the gait analysis can be further used for medical diagnosis. We believe the comfort fitting and auto-adjust with the inflatable tube was another unique feature of the work that we presented in this manuscript. It is the first time when someone combined multiple technologies and created a market-ready product for the consumer market.

## Compliance with Ethical Standards

### Conflicts of interest

Authors N. Mehendale, N. Shah and L. Kamdar declares that he has no conflict of interest also author D. Gokalgandhi declares that she has no conflict of interest aswell.

### Involvement of human participant and animals

This article does not contain any studies with animals performed by any of the authors. And also this article contain studies with human participants wearing the smart shoes. All the necessary permissions were obtained from Institute Ethical committee and conserned authorities.

### Information about informed consent

Informed consent was obtained from all the human participation who participated in wearing these smart shoes.

### Funding

There was no funding involved.

## Notes

### Competing Interest Statement

The authors have declared no competing interest.

## References

[1] N.S. Shenck, J.A. Paradiso, Energy scavenging with shoe-mounted piezoelectrics, IEEE micro 21(3), 30 (2001)

[2] K. Kong, J. Bae, M. Tomizuka, in Sensors and Smart Structures Technologies for Civil, Mechanical, and Aerospace Systems 2008, vol. 6932 (International Society for Optics and Photonics, 2008), vol. 6932, p. 69322G

[3] P.G. Jung, S. Oh, G. Lim, K. Kong, A mobile motion capture system based on inertial sensors and smart shoes, Journal of Dynamic Systems, Measurement, and Control 136(1) (2014)

[4] B.M. Eskofier, S.I. Lee, M. Baron, A. Simon, C.F. Martindale, H. Gaßner, J. Klucken, An overview of smart shoes in the internet of health things: gait and mobility assessment in health promotion and disease monitoring, Applied Sciences 7(10), 986 (2017)

[5] Z. Zhang, Z. Zhu, B. Bazor, S. Lee, Z. Ding, T. Pan, Feetbeat: A flexible iontronic sensing wearable detects pedal pulses and muscular activities, IEEE Transactions on Biomedical Engineering 66(11), 3072 (2019)

[6] H. Xia, J. Charlton, M. Hunt, P. Shull, Preliminary test of a smart shoe for training foot progression angle during walking, Osteoarthritis and Cartilage 27, S64 (2019)

[7] I. Zerin, Lpcoms: Towards a low power wireless smart-shoe system for gait analysis in people with disabilities (2015)

[8] T.L. Wood. Smart shoes (1994). US Patent 5,373,651

[9] O. Oshin, O. Oni, A. Atayero, et al., Development of a power-harnessing smart shoe system with outdoor navigation (2017)

[10] P.S. Koli, N.S. Chavan, S.G. Gaikar, Construction of smart shoe system for visually impaired people

[11] S.h. Seo, S.w. Jang, Design and implementation of a smart shoes module based on arduino, Journal of the Korea Institute of Information and Communication Engineering 19(11), 2697 (2015)

[12] R. Larkin. Meet the new “talking shoes” by google (2016). URL https://www.esoftload.info/meet-the-new-talking-shoes-by-google

[13] A. Liszewski. Reebok pumps are back and aren’t as hideous or expensive any more (2015). URL https://gizmodo.com/reebok-pumps-are-back-and-arent-as-hideous-or-expensive-1689855875

[14] E. Velloso, D. Schmidt, J. Alexander, H. Gellersen, A. Bulling, The feet in human-computer interaction: A survey of foot-based interaction, ACM Computing Surveys (CSUR) 48(2), 1 (2015)

[15] A. Gatto, E. Frontoni, in 2014 IEEE/ASME 10th International Conference on Mechatronic and Embedded Systems and Applications (MESA) (IEEE, 2014), pp. 1–6

[16] A. Gupta, A. Sharma, Piezoelectric energy harvesting via shoe sole, International Journal of New Technology and Research 1(6) (2015)

[17] E. Frontoni, A. Mancini, P. Zingaretti, A. Gatto, in ASME 2013 International Design Engineering Technical Conferences and Computers and Information in Engineering Conference (American Society of Mechanical Engineers Digital Collection, 2013)

[18] P.B. Shull, H. Xia, in 2018 IEEE 23rd International Conference on Digital Signal Processing (DSP) (IEEE, 2018), pp. 1–4

[19] S.S. Lee, S.T. Choi, S.I. Choi, Classification of gait type based on deep learning using various sensors with smart insole, Sensors 19(8), 1757 (2019)

[20] N. Mehendale, Facial emotion recognition using convolutional neural networks (ferc), SN Applied Sciences 2(3), 1 (2020)

