## supplementary information for "Walking pattern analysis using deep learning for energy harvesting smart shoes with IoT"

**S1. Embedded C Code for data transfer from Shoe to ThingSpeak**

#include <Wire.h>

#include "ThingSpeak.h"

#include <ESP8266WiFi.h>

const char* ssid	= "SSID";

const char* password = "PASSWORD";

int keyIndex = 0; // your network key Index number (needed only for WEP)

WiFiClient client;

unsigned long myChannelNumber = 978961;

const char * myWriteAPIKey = "KWRY5OECU9TVAPAO";

int number1 = 0;

int number2 = random(0,100);

int number3 = random(0,100);

int number4 = random(0,100);

int number5 = random(0,100);

int number6 = random(0,100);

int number7 = random(0,100);

String myStatus = "";

// MPU6050 Slave Device Address

const uint8_t MPU6050SlaveAddress = 0x68;

// Select SDA and SCL pins for I2C communication

const uint8_t scl = D6;

const uint8_t sda = D7;

// sensitivity scale factor respective to full scale setting provided in datasheet

const uint16_t AccelScaleFactor = 16384;

const uint16_t GyroScaleFactor = 131;

// MPU6050 few configuration register addresses

const uint8_t MPU6050_REGISTER_SMPLRT_DIV	= 0x19;

const uint8_t MPU6050_REGISTER_USER_CTRL	= 0x6A;

const uint8_t MPU6050_REGISTER_PWR_MGMT_1	= 0x6B;

const uint8_t MPU6050_REGISTER_PWR_MGMT_2	= 0x6C;

const uint8_t MPU6050_REGISTER_CONFIG	= 0x1A;

const uint8_t MPU6050_REGISTER_GYRO_CONFIG = 0x1B;

const uint8_t MPU6050_REGISTER_ACCEL_CONFIG = 0x1C;

const uint8_t MPU6050_REGISTER_FIFO_EN	= 0x23;

const uint8_t MPU6050_REGISTER_INT_ENABLE	= 0x38;

const uint8_t MPU6050_REGISTER_ACCEL_XOUT_H = 0x3B;

const uint8_t MPU6050_REGISTER_SIGNAL_PATH_RESET = 0x68;

int16_t AccelX, AccelY, AccelZ, Temperature, GyroX, GyroY, GyroZ;

void setup() {

Serial.begin(115200); // Initialize serial

WiFi.mode(WIFI_STA);

ThingSpeak.begin(client); // Initialize ThingSpeak

Wire.begin(sda, scl);

MPU6050_Init();

}

void loop()

{

// Connect or reconnect to WiFi

if(WiFi.status() != WL_CONNECTED){

Serial.print("Attempting to connect to SSID: ");

Serial.println(ssid);

while(WiFi.status() != WL_CONNECTED){

WiFi.begin(ssid, password);

Serial.print(".");

delay(5000);

}

Serial.println("\nConnected.");

}

// set the fields with the values

ThingSpeak.setField(1, number1);

ThingSpeak.setField(2, number2);

ThingSpeak.setField(3, number3);

ThingSpeak.setField(4, number4);

ThingSpeak.setField(5, number5);

ThingSpeak.setField(6, number6);

ThingSpeak.setField(7, number7);

// figure out the status message

if(number1 > number2){

myStatus = String("field1 is greater than field2");

}

else if(number1 < number2){

myStatus = String("field1 is less than field2");

}

else{

myStatus = String("field1 equals field2");

}

ThingSpeak.setStatus(myStatus);

// write to the ThingSpeak channel

int x = ThingSpeak.writeFields(myChannelNumber, myWriteAPIKey);

if(x == 200){

Serial.println("Channel update successful.");

}

else{

Serial.println("Problem updating channel. HTTP error code " + String(x));

}

// change the values

number1++;

if(number1 > 99){

number1 = 0;

}

number2 = random(0,100);

number3 = random(0,100);

number4 = random(0,100);

number5 = random(0,100);

number6 = random(0,100);

number7 = random(0,100);

delay(5000); // Wait 5 seconds to update the channel again

double Ax, Ay, Az, T, Gx, Gy, Gz;

Read_RawValue(MPU6050SlaveAddress, MPU6050_REGISTER_ACCEL_XOUT_H);

//divide each with their sensitivity scale factor

Ax = (double)AccelX/AccelScaleFactor;

Ay = (double)AccelY/AccelScaleFactor;

Az = (double)AccelZ/AccelScaleFactor;

T = (double)Temperature/340+36.53;

Gx = (double)GyroX/GyroScaleFactor;

Gy = (double)GyroY/GyroScaleFactor;

Gz = (double)GyroZ/GyroScaleFactor;

Serial.print("Ax: "); Serial.print(Ax);

Serial.print(" Ay: "); Serial.print(Ay);

Serial.print(" Az: "); Serial.print(Az);

Serial.print(" T: "); Serial.print(T);

Serial.print(" Gx: "); Serial.print(Gx);

Serial.print(" Gy: "); Serial.print(Gy);

Serial.print(" Gz: "); Serial.println(Gz);

delay(100);

}

void I2C_Write(uint8_t deviceAddress, uint8_t regAddress, uint8_t data){

Wire.beginTransmission(deviceAddress);

Wire.write(regAddress);

Wire.write(data);

Wire.endTransmission();

}

// read all 14 register

void Read_RawValue(uint8_t deviceAddress, uint8_t regAddress){

Wire.beginTransmission(deviceAddress);

Wire.write(regAddress);

Wire.endTransmission();

Wire.requestFrom(deviceAddress, (uint8_t)14);

AccelX = (((int16_t)Wire.read()<<8) | Wire.read());

AccelY = (((int16_t)Wire.read()<<8) | Wire.read());

AccelZ = (((int16_t)Wire.read()<<8) | Wire.read());

Temperature = (((int16_t)Wire.read()<<8) | Wire.read());

GyroX = (((int16_t)Wire.read()<<8) | Wire.read());

GyroY = (((int16_t)Wire.read()<<8) | Wire.read());

GyroZ = (((int16_t)Wire.read()<<8) | Wire.read());

}

//configure MPU6050

void MPU6050_Init(){

delay(150);

I2C_Write(MPU6050SlaveAddress, MPU6050_REGISTER_SMPLRT_DIV, 0x07);

I2C_Write(MPU6050SlaveAddress, MPU6050_REGISTER_PWR_MGMT_1, 0x01);

I2C_Write(MPU6050SlaveAddress, MPU6050_REGISTER_PWR_MGMT_2, 0x00);

I2C_Write(MPU6050SlaveAddress, MPU6050_REGISTER_CONFIG, 0x00);

I2C_Write(MPU6050SlaveAddress, MPU6050_REGISTER_GYRO_CONFIG, 0x00);//set +/-250 degree/second full scale

I2C_Write(MPU6050SlaveAddress, MPU6050_REGISTER_ACCEL_CONFIG, 0x00);// set +/- 2g full scale

I2C_Write(MPU6050SlaveAddress, MPU6050_REGISTER_FIFO_EN, 0x00);

I2C_Write(MPU6050SlaveAddress, MPU6050_REGISTER_INT_ENABLE, 0x01);

I2C_Write(MPU6050SlaveAddress, MPU6050_REGISTER_SIGNAL_PATH_RESET, 0x00);

I2C_Write(MPU6050SlaveAddress, MPU6050_REGISTER_USER_CTRL, 0x00);

}
